# *Staphylococcus aureus* functional amyloids catalyze degradation of β-lactam antibiotics

**DOI:** 10.1101/2023.02.01.526669

**Authors:** Elad Arad, Nimrod Golan, Hanna Rapaport, Meytal Landau, Raz Jelinek

## Abstract

Antibiotic resistance of bacteria is considered one of the most alarming developments in modern medicine. While varied pathways for bacteria acquiring antibiotic resistance have been identified, there still are open questions concerning the mechanisms underlying resistance. Here, we show that alpha phenol-soluble modulins (PSMα’s), functional bacterial amyloids secreted by *Staphylococcus aureus*, catalyze breakup of β-lactams, a prominent class of antibiotic compounds. Specifically, we show that PSMα2 and, particularly, PSMα3 catalyze hydrolysis of the amide-bond four-member ring of nitrocefin, a widely used β-lactam surrogate. Microscopic and spectroscopic analyses of several PSMα3 variants and correlation with their catalytic activities allowed mapping of the catalytic sites on the amyloid fibrils’ surface, specifically underscoring the key roles of the cross-α fibril organization, and the combined electrostatic and nucleophilic functions of the lysine residue array. This study unveils a previously unknown role of functional bacterial amyloids as catalytic agents for antibiotic compounds, pointing to possible mechanisms for antibiotic resistance of bacteria.

**ToC Graphical Abstract:** 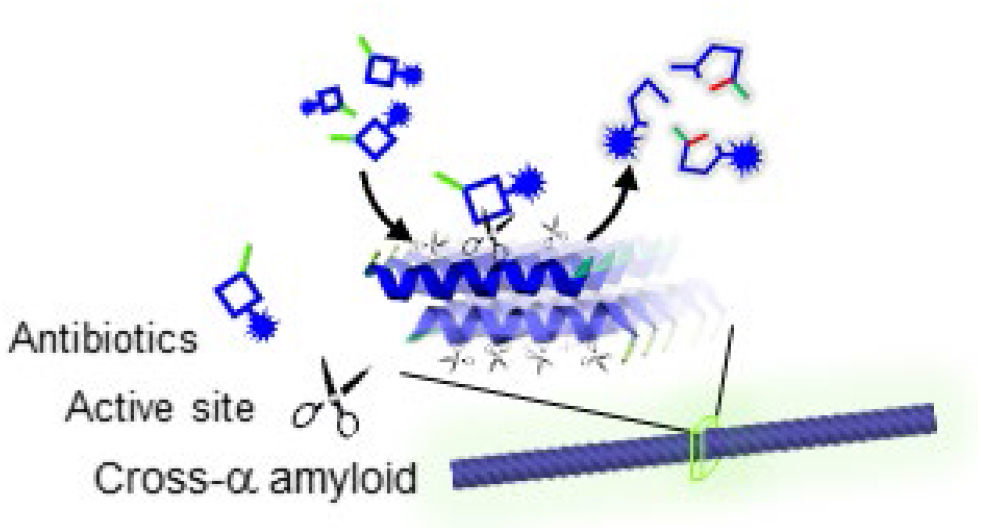

## Introduction

Antibiotic resistance by pathogenic bacteria is among the most pressing challenges in modern medicine and healthcare^1^. Indeed, pathogenic bacteria and other microorganisms have been found to employ various mechanisms to fight antibiotic potency, including genetic mutations, evolving specific receptors targeting antibiotics which activate cascades of enzymatic antibiotic-degradation^2^, and formation of physical protective barriers which adsorb or block antibiotic molecules from reaching their targets^3^. Methicillin-resistant *Staphylococcus Aureus* (MRSA), in particular, is considered one of the most health threatening antibiotic-resistant pathogens^2,4,5^. Antibiotic resistance by *S. aureus* have been traced to alteration of target proteins’ binding sites^6^. Another potent resistance strategy is the generation of antibiotic-degrading enzymes such as β-lactamases which degrade β-lactam antibiotics^7^. Indeed, antibiotic resistance is particularly acute in the case of β-lactams, which include penicillin, penems, and others^8^, that are among the most commonly-used drugs worldwide.

Bacterial biofilms, the rigid matrix produced by many pathogenic bacteria, also play a role in antibiotic resistance as biofilm layers have been shown to adsorb small molecules thereby putatively blocking access to the bacterial colonies^9^. Biofilm frameworks comprise varied biomolecules, major among them are protein amyloid fibers^10^. Amyloid fibrils, observed in diverse protein systems and mostly associated with neurodegenerative diseases^11^, have been also increasingly encountered in bacterial biology. Functional bacterial amyloids (referred to as FuBAs) serve as building blocks for biofilm scaffolding^12^, facilitate resistance to stress conditions^13^, act as bio-adhesives anchoring agents^14^, create channels in the biofilm matrix for nutrient uptake and transport^15,16^, and even act as antibacterial agents, achieving dominance over other microbial species^17–19^. While prominent FuBAs, including *curli* (secreted by *E. coli*)^20,21^, *Fap (P*.*aeruginosa)*^8^, and *tas (bacillus)*^16^ adopt β-sheet amyloid structures, amyloid proteins produced by *S. aureus*, specifically belonging to the *phenol-soluble modulin* (PSM) family have been shown to form unique cross-α organizations^22,23^. Specifically, the PSMα peptides which are highly amphiphilic, induce the formation of α-helical bilayers that bury their hydrophobic residues and expose the hydrophilic ones^24^.

Amyloidogenic proteins have been shown to exhibit catalytic properties^25–27^. Self-assembled peptides were reported to catalyze hydrolysis, condensation, redox reactions^26,27^, and even multi-step cascade-reactions^28,29^. While almost all previous studies have focused on synthetic *de-novo* peptides or amyloidogenic protein domains as catalytic agents^30–34^, we have recently demonstrated that native, physiological amyloids can catalyze disease-related reactions^35^ and key metabolic reactions^36^. These discoveries suggest that amyloid proteins may act as catalysts for varied biological reactions and functional targets. Here, we demonstrate that members of the PSMα family, particularly PSMα3, catalyze breakup of β-lactam antibiotics. Through structure/function analyses of PSMα3 variants, putative catalytic sites on the amyloid fibrils’ surface are localized, linked to both the cross-α organization of the amyloid fibrils and electrostatic interactions between the β-lactam molecules and lysine sidechains. amyloid-catalyzed β-lactam degradation may underscore an intriguing new paradigm accounting for antibiotic resistance of *S. aureus*, and pathogenic bacteria in general.

## Results and discussion

The hypothesis underlining this study is illustrated in **Figure 1**. β-lactams adsorb PSMα amyloid fibrils (physiologically identified as constituents of *S. auereus* biofilms, and shown to be secreted by this bacterial species as toxins^22^), and catalytically degraded through amide bond hydrolysis in the 4-member ring. Structurally, β-lactams contain an electrophilic, activated-amide bond ring, allowing it to react with various enzyme nucleophiles leading to bacterial cell destruction^8^. The experiments presented below are designed to examine whether PSMα amyloids, through active sites upon the amyloid fibril surface (schematically represented as “scissors”), catalyze cleavage of the four-membered β-lactam ring, rendering the molecule functionally inactive. Specifically, we tested PSMα amyloid-mediated degradation of nitrocefin, a widely-studied β-lactam surrogate of penicillin^37^.

**Figure 1.**
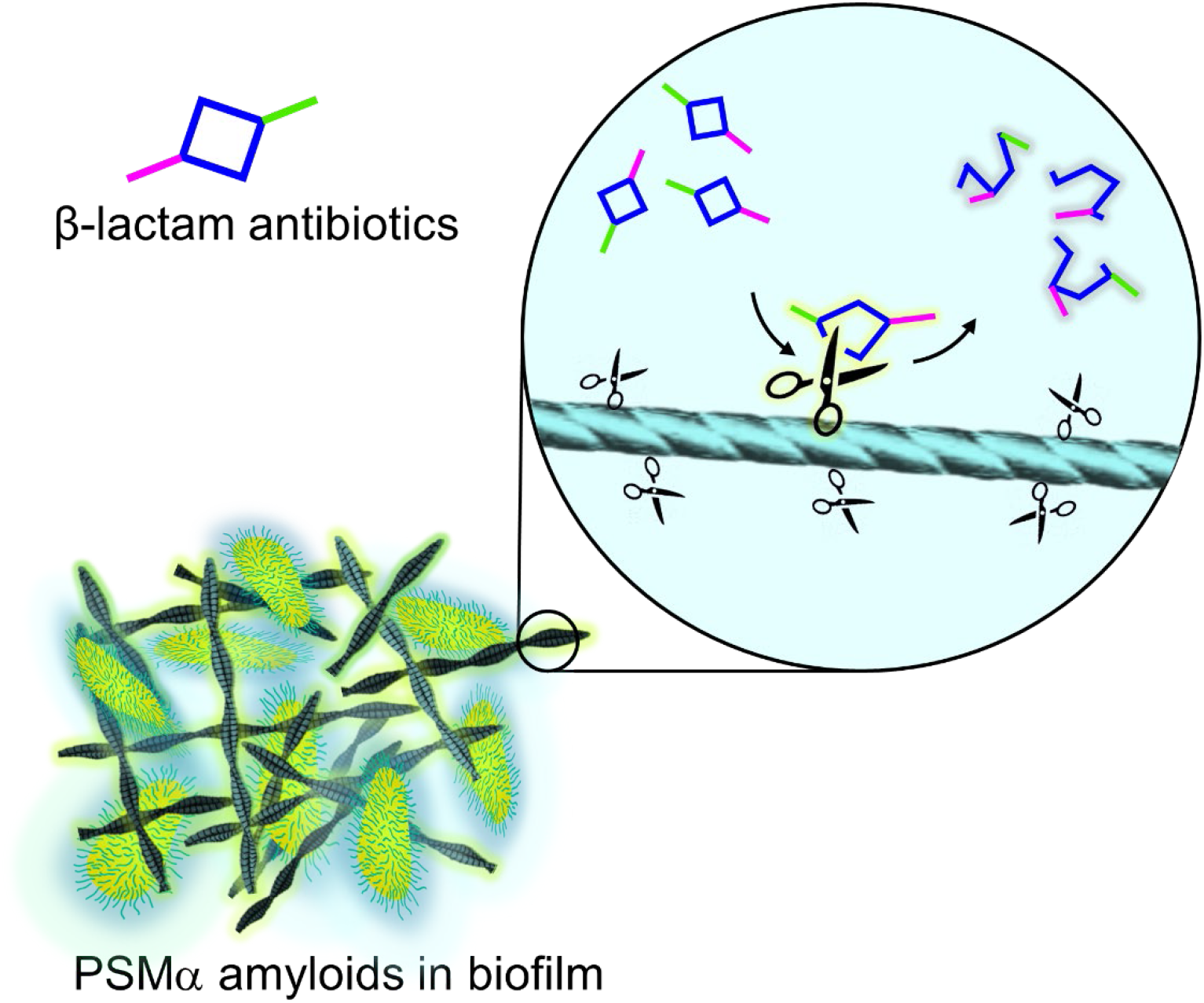
*S. aureus-*secreted functional amyloids catalytically degrade β-lactam antibiotics. PSMα amyloids adsorb β-lactams and catalyze their degradation through amide bond hydrolysis.

**Figure 2**. presents structural analyses of the PSMα1-4 amyloid fibrils investigated. Figure 2A depicts the amino acid sequences of the peptides, highlighting in different colors the amyloid surface-displayed residues which may play roles in catalysis. Specifically, acidic residues (red), basic residues (blue), or neutral nucleophilic residues (turquoise) have been observed in enzymatic active sites^38–40^. Particularly, in many enzymes, polar residues in the active sites play prominent functions by binding substrate molecules, activating proximate residues, or partaking in nucleophilic attacks^40^.

**Figure 2.**
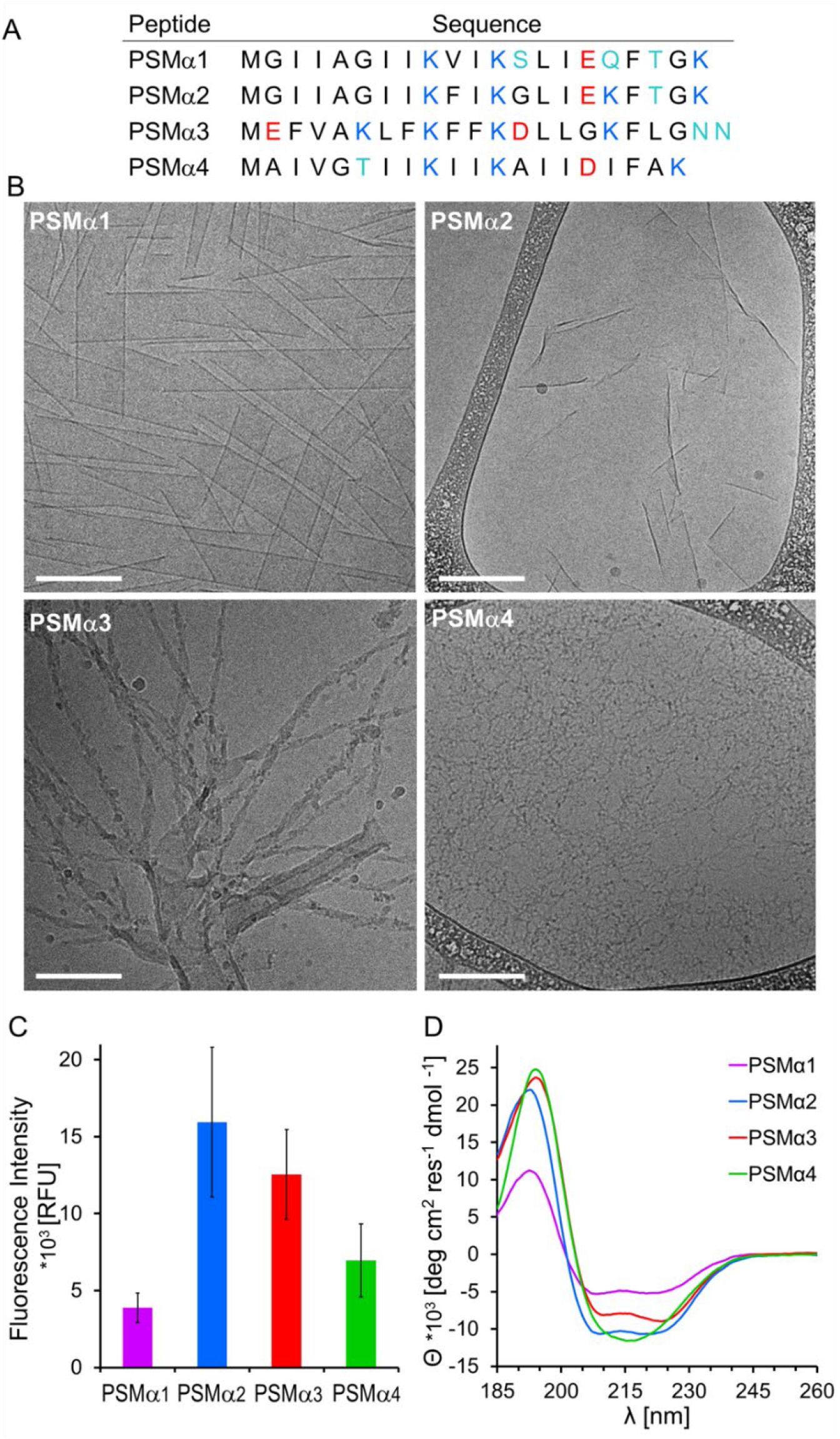
Structural characterization of the PSMα amyloids. **A**. Sequences of PSMα 1-4. Anionic, cationic, and nucleophilic residues are marked in red, blue and turquoise, respectively. **B**. Cryo-TEM images of the peptides (concentration 300 μM). Bars correspond to 200 nm. All PSMα samples were pre-incubated in water for two hours and buffered with HEPES prior to image acquisition. **C**. Amyloid staining with *Amytracker 680* (excitation 550 nm, emission 650 nm, PSMα concentrations were 400 μM). **D**. Circular dichroism (CD) spectra of the peptides (concentration 170 μM, peptides dissolved in HEPES buffer, pH=7.4).

Figure 2B shows cryogenic transmission electron microscopy (cryo-TEM) images of the PSMα peptides, following two hours incubation in deionized water, followed by buffering with HEPES (pH 7.4). Both the incubation period and relatively high concentration (300 μM) allowed the PSMα to attain maximal aggregation. The cryo-TEM analysis reveals interesting morphological differences between the peptide assemblies. Specifically, PSMα1 adopted ∼100 nm-wide filaments while PSMα2 formed extended thin sheets having varying dimensionalities. PSMα3, in contrast, mainly assembled into typical long twisted fibrils (with tubular structures also observed)^41^. The cryo-TEM image of PSMα4 reveals that this peptide did not exhibit organized structures but rather produced small aggregates.

Fluorescence emission measurements of PSMα fibrillar samples incubated with the amyloid-specific dye *Amytracker*-680, which fluorescence intensity correlates with repeating amyloidogenic and positively-charged amyloid domains^42,43^, are portrayed in Figure 2C. The bar diagram in Figure 2C indicates that variability in amyloid formation by the PSMα assemblies was apparent. PSMα2 and PSMα3 gave rise to high fluorescence of the dye, indicating pronounced amyloid organization, while PSMα1 and PSMα4 generated lesser *Amytracker*-680 fluorescence, reflecting lower amyloid organization within the peptide aggregates. The circular dichroism (CD) analysis in Figure 2D further illuminates the secondary structures of the PSMα peptides. The CD spectra of PSMα1, PSMα2, and PSMα3 indicate that the three peptides adopted α-helical structures, reflected in the maxima at 195 nm and minima at 210 nm and 225 nm^23,44^, as these polypeptides form cross-α amyloid fibrils^45,46^. PSMα4, on the other hand, featured β-sheet assemblies, accounting for the spectral minimum in 215 nm and peak at 198 nm, consistent with formation of cross-β fibril organization^22,47^. These secondary structures were also confirmed by FTIR amide-I spectroscopy (Figure S1).

Figure 3 explores PSMα-mediated catalysis of β-lactam ring breakup via amide bond hydrolysis. In the experiments, we monitored the four-membered ring opening in nitrocefin following addition of the PSMα amyloids (pre-incubated at concentration of 400 μM in water for two hours and then buffered with HEPES, pH 7.4) and tracking the absorbance at 480 nm^37^. Figure 3A reveals significantly different catalytic properties of the PSMα aggregates. Specifically,

**Figure 3.**
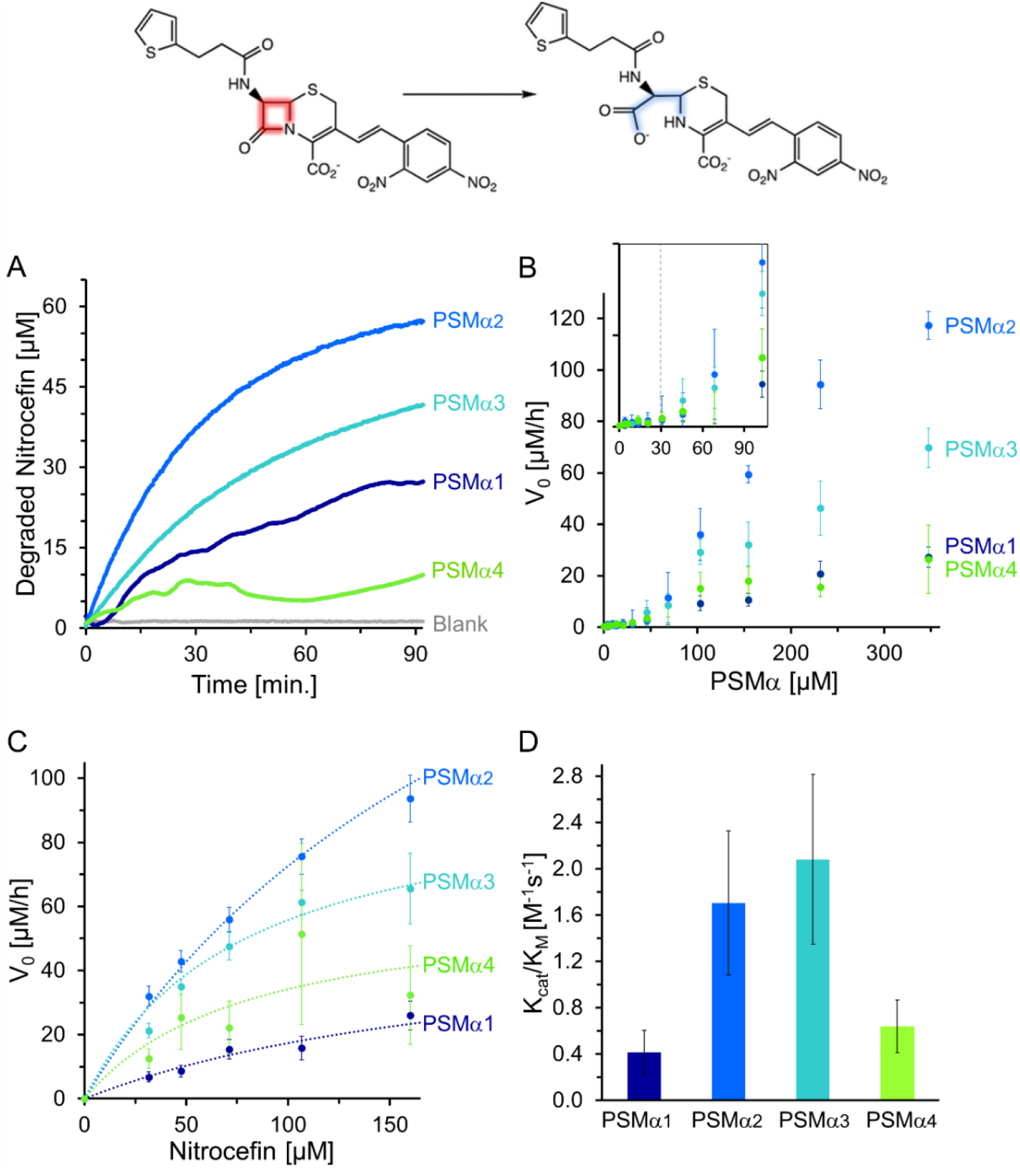
Catalytic amide bond hydrolysis of nitrocefin by the PSMα assemblies. A scheme of nitrocefin β-lactam ring opening and degradation (top row). **A**. Graph depicting degradation of nitrocefin (initial nitrocefin concentration 60 μM) in the presence of PSMα assemblies (350 μM). **B**. Initial nitrocefin-degradation reaction rate, V_0_, as a function of initial preassembled-PSMα concentration (the peptides were pre-incubated in water for two hours and then buffered with HEPES prior to mixing with nitrocefin at concentration of 60 μM). The inset depicts the low-concentration peptide region, underscoring a negligible catalytic activity at V_0_ lower than 30 μM (marked in broken vertical line). **C**. Initial nitrocefin-degradation reaction rate as function of initial nitrocefin concentration (PSMα peptide concentrations of 170 μM). Each point represents an average ± SEM, N_PSMα1_=8, N_PSMα2_=19, N_PSMα3_=10, N_PSMα4_=5. The dotted line is the fitting to Michaelis-Menten (MM) model using non-linear regression. **D**. The catalytic efficiency (K_cat_/K_M_) as derived from the fitting to MM model (using the catalytic parameters in Table S1). Values are average ± confidence interval.

PSMα2 and PSMα3 amyloids markedly accelerated nitrocefin degradation, resulting in almost full turnover in the case of PSMα2 and 75% for PSMα3. PSMα1 gave rise to lower turnover of ∼50%, while PSMα4 induced minor degradation of nitrocefin (turnover of ∼15%).

Figure 3B depicts the initial nitrocefin degradation reaction rates, *V*_0_ (calculated from linear fitting of the degraded-nitrocefin absorbance slopes, Figure 3A, in the initial 15 minutes), as a function of PSMα peptide concentrations (initial nitrocefin concentration 60 μM). Consistent with the kinetic data in Figure 3A, PSMα2 amyloids exhibited the highest *V*_*0*_ followed by PSMα3 aggregates, while PSMα1 and PSMα4 exhibited lesser catalytic activities (i.e., low initial degradation of nitrocefin). Interestingly, Figure 3B reveals that the *V*_*0*_ for all peptides exhibited linear dependency reflecting higher catalytic activity when greater amyloid concentrations were present in the aqueous solution. Importantly, nitrocefin breakup was observed only above a ∼30 μM threshold (Figure 3B, inset). Indeed, we observed that below this threshold concentration all four PSMα amyloids were practically soluble (Figure S2). Overall, Figure 3B attests to a direct relationship between PSMα amyloid formation and nitrocefin degradation.

Measurements of *V*_*0*_ as function of initial substrate (nitrocefin) concentrations (keeping the PSMα concentration constant at 170 μM, assuring amyloid formation, e.g., above the 30 μM threshold) are depicted in Figure 3C. Notably, the shapes of the four curves (accounting for the respective PSMα peptides) indicate a Michaelis-Menten (MM) kinetic dependence, corroborating an enzyme-like catalytic activity^48^. Table S1 summarizes the pertinent enzymatic parameters extracted from the initial reaction rate data in Figure 3C using non-linear regression fit to a MM reaction model^48^. Figure 3D presents the catalytic efficiencies, calculated for each peptide by dividing the turnover rate by the MM-constant (K_cat_/K_M_), considered a measure for both substrate binding affinity and conversion kinetics into product. Consistent with the data in Figures 3A-C, the bar diagram in Figure 3D demonstrates that PSMα2 and PSMα3 exhibit significantly higher catalytic efficiencies compared to PSMα1 and PSMα4.

To decipher the structural parameters and putative active sites in the most catalytically efficient peptide amyloid -PSMα3, we analyzed the structural and catalytic properties of several PSMα3 variants, in which specific residues were altered in strategic positions (**Figure 4)**. Figure 4A shows the sequences of the PSMα3 variants tested. *α3-cationic* has pronounced positive charge since all acidic functional groups were amidated (the glutamic acid in position 2, aspartic acid in position 13 and asparagine in the C-terminus), thereby avoiding possible electrostatic repulsion with the anionic nitrocefin substrate. In comparison, the lysine residues in the *α3-EG/K* variant were capped with two units of ethylene-glycol, thus blocking the positive residues and increasing the amphiphilicity of the sidechains. *α3-F10P* does not adopt cross-α architecture due to the rigid turn-motif introduced by substitution of phenylalanine with proline^22^. Similar disruption of cross-α organization was designed in the case of the *α3-(d)FF* variant, in which the overall sequence is retained but the L-Phe residues in positions 10 and 11 were substituted with diastereomeric D-Phe^22^. Another control PSMα3 variant was a scrambled sequence *(α3-scrambled*). This variant contains the same amino acids as the parent peptide, however it does not display the AABBAA pattern (in which A and represent hydrophilic and hydrophilic residues, respectively), important for α-helical folding^49^.

**Figure 4.**
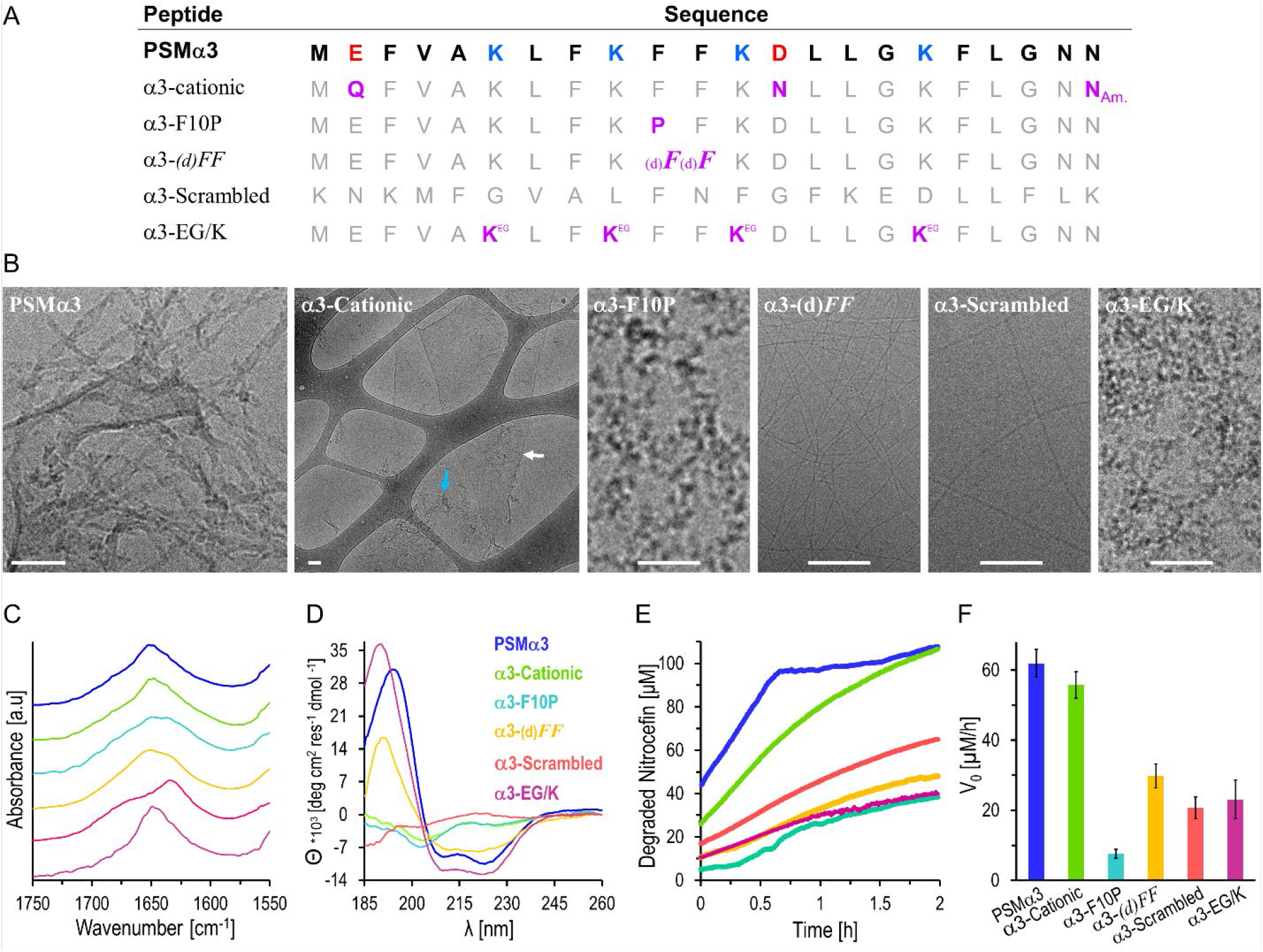
Structural features and catalytic activities of PSMα3 variants. **A**. Sequences of the PSMα3 variants. The modified residues in the variants are marked in purple. *α3-cationic*: all acidic residues and the Asn-22 in the C-terminus were amidated. *α3-F10P:Phe-10 was replaced with Pro residue. α3-(d)FF*: two D-Phe residues instead of the L-handed residues. *α3-EG/K*: amine sidechains in the Lys residues were capped with two units of ethylene-glycol. **B**. Cryo-TEM images of the PSMα3 variants (concentrations of 300 μM). Bars correspond to 100 nm. **C**. Amide region FTIR spectra; **D**. CD spectra of the PSMα3 variants. **E**. Nitrocefin degradation reaction kinetics in the presence of PSMα3 variant assemblies formed after 2-hour incubation in DIW and addition of HEPES buffer (peptide concentrations were 170 μM, initial nitrocefin concentration 107 μM). **F**. Initial degradation rates, V_0_, at initial nitrocefin concentration of 107 μM. The same color-coding was used in panels C-F.

Figure 4B shows the cryo-TEM morphologies of the PSMα3 variants following incubation for two hours in water and buffering with HEPES pH 7.4. Different than native PSMα3 assembled primarily as twisted fibrils, the *α3-cationic* variant produced micron-scale thin sheets, ascribed to pronounced effect of the electrostatic repulsion between the positive moieties, blocking sheet-to-fibril transformations^50^. Surprisingly, both *α3-scrambled* and *α3-(d)FF* appear to form thin fibrillar structures, accounting for the amphiphilic sequences of these variants and adoption of β-sheet structure. In contrast to the other variants, the cryo-TEM images in Figure 4B reveal that *α3-F10P* and *α3-EG/K* formed amorphous aggregates. In the case of *α3-F10P*, this result is likely due to the structural rigidity introduced by proline-10, disrupting folding neither into α-helices or β-sheets. The amorphous structures of *α3-EG/K* likely reflect the effects of the ethylene-glycol units decorating the four lysine residues, blocking participation of the charged lysines in peptide self-assembly. Fluorescent labelling of these variant with *Amytracker-680* is consistent with the TEM images (Figure S3).

Figure 4C depicts the attenuated total reflectance-FTIR (ATR-FTIR) spectra of the amide-I vibration region of the PSMα3 variants, pointing to their secondary structures^51^. Figure 4C demonstrates that native PSMα3 and the *α3-cationic* variant gave rise to similar peaks centered at 1655 cm^-1^, the fingerprint peak for α-helical structure^52^. *α3-F10P* also gave rise to a signal at 1647-1650 cm^-1^, although markedly broader, accounting for substantial degree of unordered random-coil organization^53^. The FTIR spectrum of *α3-(d)FF* was significantly different than the other three peptides, displaying a shoulder at 1628 cm^-1^ and a maximum at 1645 cm^-1^, indicating a combination of β-sheet and random-coil conformations^35,53^, likely due to the diphenylalanine motif known to contribute to formation of β-sheet assemblies^54^. Interestingly, *α3-scrambled* generated a spectrum displaying a maximum at 1627 cm^-1^ and a shoulder at 1684 cm^-1^, indicating an antiparallel β-sheet conformation probably due to the amphiphilic sequence lacking the repeating pattern of (hydrophobic)_2_-(hydrophilic)_2_. Like native PSMα3, *α3-EG/K*, in which the lysine residues were capped, formed α-helical structure, reflecting similar amphiphilic pattern of hydrophobic and hydrophilic residues, accounting for the hydrophilicity of the ethylene glycol capped lysine residues.

Figure 4D depicts the CD spectra of the PSMα3 variants. The CD trace of native PSMα3 indicates a typical α-helical structure (blue spectrum), with two minima at 210 nm and 225 nm. *α3-cationic* generated low-intensity spectrum indicating that most of the peptide aggregates thereby inducing light scattering (i.e., the micrometer-sized sheets in the cryo-TEM image, Figure 4B). *α3-F10P* appear to have formed polyproline type II structure with a minimum at 198-200 nm, reflecting an unordered structure rich in proline residues, adopting turn-like rather than α-helical structure (confirmed also in the amorphous aggregates seen in the cryo-TEM analysis, Figure 4B). *α3-(d)FF* yielded a typical β-sheet spectrum with minimum around 218 nm, corroborating the FTIR spectrum (∼1630 cm^-1^ shoulder, Figure 4C) and the apparent fibrillar morphology seen in the cryo-TEM analysis. *α3-scrambled* generated a very low intensity spectrum, likely linked to aggregation of the peptide. Figure 4D also shows that *α3-EG/K* produced a α-helical spectrum akin to the parent PSMα3.

Figure 4E and 4F portray the PSMα3 variant-mediated catalytic nitrocefin amide bond hydrolysis, demonstrating pronounced divergence in catalytic performance. Figure 4E depicts the nitrocefin degradation curves recorded in the presence of PSMα3 variants at a concentration of 170 μM (prepared similarly to the experiment in Figure 3C - incubated in water for two hours and buffered prior to mixing with nitrocefin). PSMα3 exhibited the most significant activity, degrading the entire nitrocefin reservoir (107 μM) after ∼40 minutes, thereby reaching a plateau. *α3-cationic* exhibited similarly significant catalytic activity as native PSMα3, although at seemingly slower rate and without reaching an early plateau, likely reflecting the abundant amine residues at the amyloid surface. Notably, all other PSMα3 variants displayed lower catalytic performance (*α3-scrambled* featured somewhat more pronounced catalytic activity, red curve in Figure 4E).

The bar diagram in Figure 4F further illuminates the catalytic profiles of the PSMα3 variants, outlining the initial rate of nitrocefin degradation reaction (*V*_*0*_), calculated from the kinetic curves in Figure 4E. Figure 4F demonstrates that native PSMα3 and the *α3-cationic* variant exhibited the highest initial reaction rates (around 60 μM/h), while the β-sheet-forming variants gave rise to lower initial rates *(α3-(d)FF* exhibited 30 μM/h, while *α3-scrambled* - 21 μM/h) and *α3-EG/K* also exhibited low *V*_*0*_ of 23 μM/h. *α3-F10P*, in comparison, had a very low catalytic activity characterized by initial reaction rate that was lower than 15 μM/h. Variant analysis was carried out in the case of PSMα2 (Figure S4), furnishing similar structure/function insight on the catalytic activities of the peptide.

## Discussion

While PSMs secreted by *S. aureus* have been reported as framework constituents in biofilm scaffoldings, the precise functions of PSMα amyloids are still debated. Here, we demonstrate for the first time that native PSMα amyloids catalyze degradation of the prominent antibiotic β-lactam. Interestingly, PSMα2 and PSMα3 featured the highest catalytic activity towards amide bond hydrolysis of nitrocefin, compared to the two other sequences tested, PSMα1 and PSMα4. Indeed, while PSMα1 and PSMα4 were shown to contribute to *S. aureus* biofilm integrity and mechanical stability^55^, PSMα3 was also reported to play a role in inflammatory processes, virulence and cytolytic activity of MRSA^56^. As such, PSMα3-mediated catalytic degradation of antibiotic compounds may constitute an important additional factor in *S. aureus* virulence and antibiotic resistance.

The divergent structural and catalytic properties of the PSMα3 variants (Figure 4) underscore important structure/function relationships pertaining to the catalytic nitrocefin degradation process. PSMα3 amyloids exhibit core structural features which may contribute to nitrocefin catalysis: their cationic nature arising from the four surface-exposed lysine residues, and the semi-periodic amphiphilic pattern of hydrophobic-hydrophobic-hydrophilic-hydrophilic residues in the sequence, known to induce twisted α-helical assembly^49,57,58^. Lys residues likely play dual roles, both serving as anchors for negatively charged molecules (such as nitrocefin), as well as functioning as nucleophiles (via the amine side chains^31^). Indeed, both PSMα2 and PSMα3, which exhibited the most pronounced catalytic activities, have four lysine residues in their sequence vs three lysines in PSMα1 and PSMα4.

The PSMα3 variant analysis (Figure 4) furnishes further evidence for the key catalytic roles of the lysine residues in PSMα3 amyloids. *α3-cationic*, where all negatively charged residues were amidated, showed similar catalytic activity in comparison to PSMα3, although forming sheets rather than fibrils. Yet, unlike PSMα3 fibrils, *α3-cationic* assemblies did not reach nitrocefin hydrolysis product saturation, indicating that higher number of active sites probably existed (experiments utilizing other peptide concentrations yielded similar outcome; Figure S5). Similarly, the major decrease in the catalytic activity of the *α3-EG/K* variant may be explained by the inability of the Lys amine sidechains to act as nucleophiles, in parallel with reducing the concentration of cationic moieties on the amyloid surface.

The structure/function analyses in Figures 2-4 also underscore the important contribution of the cross-α amyloid structure in nitrocefin degradation. *α3-F10P*, for example, while resembles PSMα3 in terms of amphiphilic nature, did not form a α-helical structure due to the rigid proline residue, leading to major decrease in catalytic activity. α3-(d)*FF* and *α3-scrambled* did fibrillate into antiparallel cross-β structure, giving rise to moderate catalytic activities, attributed to lesser exposure of lysine sidechains as in the cross-α array structure. Indeed, synthetic short cross-α peptides were recently shown to catalyze hydrolysis reactions while exposing seryl-histidine group on the fibril surface^59^. The helical structure provides softer and less rigid environment in comparison to β-sheet^60^, that can contribute to the fibril’s flexibility upon substrate binding. Moreover, the cross-α architecture can provide higher surface area, further creating domains that can function as active sites.

**Figure 5.** portrays the proposed catalytic mechanism accounting for β-lactam degradation by the PSMα3 cross-α amyloid fibrils, based upon the experimental data in Figures 2-4. While the cross-α fibril (consisting of two peptides in each layer^45^) buries the hydrophobic residues^23^, the hydrophilic residues are exposed on the fibril surface. As such, anchoring of the β-lactam substrate occurs on the fibrils surface primarily via the lysines’ amine sidechains (marked in blue). Similar behavior of *de-novo* amphiphilic β-sheet amyloidogenic structure decorated with lysines was previously shown to exhibit catalytic activity towards enantioselective chemical reactions^33,34^. Notably, the ordered lysine arrays generate higher local positive charge thus enabling enhanced adsorption of anionic molecules and increased reactivity via nucleophilic attack. In contrast, the Lys-arrays are disrupted in the case of α3-(d)*FF* or *α3-scrambled* variants, resulting in lower catalytic activities. Cross-α assemblies, unlike other aggregates, exhibit lesser rigidity^61,62^, possibly allowing efficient biding of the substrate, creation of transition states, and release of the product, that are all necessary for effective catalysis to occur.

**Figure 5.**
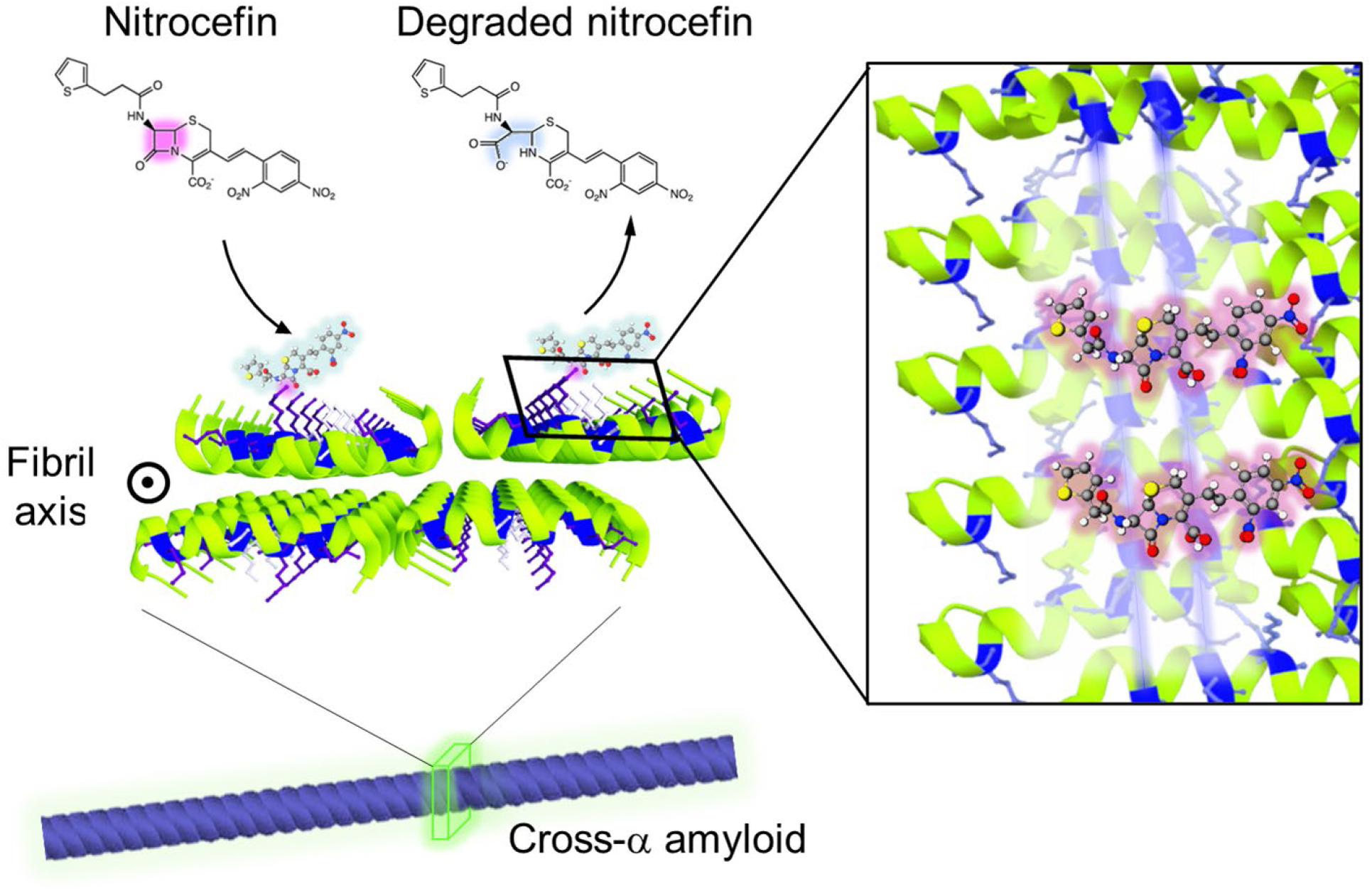
Model of PSMα3-mediated β-lactam catalysis. The cross-α amyloid fibrils, composed of bilayer made of amphiphilic α-helices, display the lysine residues on the surface of the fibril (marked in blue). Nitrocefin adsorbs onto the cationic lysine sidechains and the β-lactam ring (marked in pink) act as an electrophile while lysine residues as nucleophiles, attacking the β-lactam carboxyl. The right column depicts a top view of the fibril with nitrocefin adsorbed in parallel to the α-strands to maximize electrostatic interactions and amphiphility of the bilayer. PDB structure 5I55^23^.

The observation that functional bacterial amyloids catalyze degradation of common antibiotic small molecules furnishes a new mechanism for antibiotic resistance phenomena. While the main contribution of FuBAs was thought to be their presence in biofilm scaffolds thereby acting as physical barriers for shielding bacterial cells, catalytic degradation of antibiotic compounds may be a powerful mechanism for protecting bacteria. Previous studies have shown, for example, that FuBAs in biofilm constructs adsorb small molecule metabolites, aiding bacteria to survive in stress conditions^9^. Furthermore, different than other bacterial resistance mechanisms, the catalytic activities reported here are general and pertain to various β-lactams. Indeed, unlike some β-lactamases that are substrate-specific, amyloid catalysis may be pertinent for a wide range of β-lactam antibiotics, not requiring “lock-key” fits to specific substrates. Overall, this work may underscore new mechanistic pathways accounting for the growing threat of MRSA antibiotic resistance, particularly in the case of β-lactam antibiotics.

## Conclusions

This study presents the first evidence of catalytic properties of functional bacterial amyloids in general, particularly cross-α bacterial amyloids. We specifically show that PSMα assemblies, mainly PSMα2 and PSMα3, catalyze amide-bond hydrolysis of β-lactam antibiotics. Comprehensive structure/functional screening of PSMα3 variants illuminate the key features of the catalytically active sites on the amyloid fibrils’ surface, specifically the cross-α organization and key roles of the lysine residues in anchoring the β-lactam molecules via electrostatic attraction and nucleophilic attack resulting in amide bond hydrolysis and breakup of the four-member lactam rings. Our observations outline a previously unknown function of bacterial amyloids and may highlight contribution of bacterial biofilms in antibiotic resistance.

## Experimental Section

### Materials

Phenol Soluble Modulin α (PSMα) peptides and their derivatives were synthesized, purified by high performance liquid chromatography to > 95%, characterized by mass spectroscopy and supplied as a lyophilized powder by GenScript (Piscataway, NJ). 4-(2-hydroxyethyl)-1-piperazineethanesulfonic acid) (HEPES, >99% purity) and Nitrocefin (3-(2,4-Dinitrostyryl)-(6R,7R)-7-(2-thienylacetamido)-ceph-3-em-4-carboxylic Acid, M.W of 516.50) were purchased from Holland-Moran (Yehud, Israel). Ammonium-acetate (99% purity), 1,1,1,3,3,3-Hexafluoro-2-propanol (HFIP, 99% purity), were purchased from Sigma-Aldrich (Rehovot, Israel). Acetonitrile (LC-MS grade, 99.9% purity) was purchased from Beith Dekel (Raanana, Israel. Manufactured by JTBaker). Amytracker-680 fluorescent dye was supplied by Ebba biotech (Stockholm, Sweden).

### Peptide sample preparation

PSMα peptides and derivatives were dissolved in HFIP and titrated with NH_4_OH (30%) until the solution was clear and no turbidity was detected, to prevent aggregation and solubilize in monomeric forms. A stock solution at a concentration of 10 mM was prepared and stored at −20 °C until use. For each experiment, the solution was thawed, and the required amount was separated to aliquots in glass test-tubes and dried by evaporation (0-40 mBarr) for two hours to remove the HFIP (in most cases, 20 μL of concentrated peptide was placed in each aliquot test tube). The dried peptide samples were dissolved in doubly ionized water (DIW) applying short vortex and incubated for two hours at room temperature at peptide concentration of 444 μM. Then, buffer and DIW were added to dilute the sample to final concentration of 400 μM (in all kinetic experiments the buffer was HEPES 50 mM, pH=7.4, in CD experiments the HEPES concentration was 5 mM). The buffers were prepared with DIW, 18.2 MΩcm (Barnstead Smart2Pure, Thermo Scientific).

### Amyloid quantification and labelling

PSMα assemblies were prepared as described above, incubated in DIW for two hours at concentration of 440 μM and then buffered with HEPES (final PSMα concentration 400 μM, HEPES 50 mM). Amytracker-680 was diluted with DIW by 50. 55 μL of PSMα solution was mixed with 5 μL of Amytracker-680 stock solution (Amytracker-680 final dilution was *500). Then, the samples were placed in 384 well black plate, incubated for 10 minutes and then fluorescent measurement was applied (Excitation at 550 nm, emission 650 nm) at Biotek Synergy H1 plate reader (Biotek, Winooski, VT, USA).

### Circular Dichroism (CD) Spectroscopy

CD spectra were recorded in a range of 185–260 nm on a Jasco J-715 spectropolarimeter (Tokyo, Japan), with 0.01 cm quartz cuvettes. All samples were prepared in diluted HEPES buffer of 5 mM to avoid buffer dependent noise and recorded at final peptide concentration of 170 μM. CD spectra were recorded at 0.5 nm wavelength data pitch at 20°C and represents an average of four scans. Background CD signals of the buffer recorded and subtracted from the corresponding spectra. The ellipticity (θ) was normalized in accordance to the pathlength, the peptide concentration and number of residues (presented as θ_MRE_).

### Fourier Transform Infrared Spectroscopy (FTIR)

Peptide samples were prepared as described above, in ammonium acetate buffer (10 mM, pH =7.35). Ammonium-acetate buffer was used due to its ability to partially evaporate along the lyophilization, allowing smaller buffer fingerprint and noise. Following the incubation, the samples were placed in plastic test-tubes and cryogenically frozen by dipping in liquid nitrogen. Following this, the samples were lyophilized at -30°C to preserve the secondary structure (LABCONCO-Triad, Labotal, Israel). FTIR spectra was recorded using NICOLET 6700 FTIR spectrometer with attenuated total reflectance (ATR) system equipped with diamond crystal and DTGS detector (Thermo Fischer Scientific, MA, USA). The FTIR spectra were acquired from 122 scans at 4 cm^-1^ resolution and were corrected for spectral distortion using atmospheric suppression. A baseline correction function was applied to all spectra. Reference spectra were measured using bare ATR crystal surface.

### Cryogenic transmission Electron Microscopy (cryo-TEM)

a 5 μL droplet of each peptide’s solution, prepared as reported above, was deposited on a glow-discharged TEM grid (300 mesh Cu Lacey substrate grid; Ted Pella). The excess liquid was automatically blotted out with a filter paper and the specimen was rapidly plunged into liquid ethane precooled with liquid nitrogen in a controlled environment (Leica EM GP). The vitrified samples were transferred to a cryo-specimen holder and examined at −181 °C with a FEI Talos S200C microscope operated at 200 kV in low-dose mode. Images were recorded using a Ceta camera (4k × 4k) and analyzed using Digital Micrograph Gatan Inc. software.

### Nitrocefin hydrolysis kinetics

Nitrocefin, a β-lactam that is widely used as a surrogate substrate of Penicillin in β-lactamase activity assay, was used as a substrate. NC degradation was monitored for two hours. Nitrocefin stock solution was dissolved in acetonitrile (20 mM) and then diluted with HEPES buffer (50 mM, pH 7.4) to nitrocefin concentration of 360 μM (further dilutions to lower concentrations were made with the 360 μM NC stock). All nitrocefin samples were kept on ice in cold conditions and any further dilution or mixing was performed in ice-cold conditions to avoid self-hydrolysis background reaction. 120 μL of the pre-cooled nitrocefin solutions were mixed inside pre-cooled clear 96-well plate (Greiner F-bottom) with 60 μL of pre-incubated PSMα solution (510 μM PSMα, incubated in DIW and later buffered in HEPES 50 mM, pH 7.4). Immediately after mixing the samples, the plate was transferred to 20° C pre-cooled plate reader (Multiskan GO, Thermo scientific, Waltham, MA) and shaken slowly for 10 seconds. Then, the absorbance of degraded nitrocefin was measured for two hours at interval of 20 seconds at λ=480 nm. Initial production rates (V_0_) were calculated by slope of the initial linear range of degraded-nitrocefin production at the first 15 minutes. The reported results are and average of at least five separate measurements (the exact number of repeats, N, is reported next to the values). Each experiment included also blank samples of nitrocefin substrate diluted with buffer instead of PSMα. All presented results include subtraction of buffer control. The initial rate calculations were performed while also subtracting the self-degradation of the substrate, to ensure that this product absorbance is a result of PSMα presence.

### Kinetic data fit to the Michalis-Menten (MM) model

The initial rates of degraded-nitrocefin production were calculated from the slopes of the first 15 minutes of measurement as reported above. The initial rates (V_0_) were fitted using non-linear regression applied with the solver function in excel, either to Michalis-Menten equation

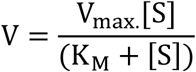

Where V_max._ Stands for maximal reaction rate, K_M_ stands for Michalis-Menten constant, [S] stands for the substrate molar concentration. Each initial rate value was presented as average ± SEM. The kinetic parameters were calculated by fitting the data to MM model using non-linear regression, and the confidence intervals for each parameter were calculated accordingly.

## Acknowledgements

The measurements of electron microscopy were performed at the Ilse Katz Institute (IKI) for Nanoscale Science and Technology Ben-Gurion University of the Negev, Beer Sheva, Israel. The authors are thankful to Dr. Einat Nativ-Roth for her great support, fruitful comments and help in sample preparation for electron microscopy and imaging of cryogenic TEM. The authors are also thankful to Dr. Alexander Upcher, for his great help in sample preparation for microscopy measurements. The authors admire Dr. Sofiya Kolusheva for her critical comments along the project.

## ASSOCIATED CONTENT

## Supporting Information

Figure S1. FTIR spectroscopy of PSMα1-4

Figure S2. Critical dissolution concentration of PSMα1-4.

Table S1. Catalytic parameters of PSMα1-4 amyloids

Figure S3. Amytracker-680 fluorescent labelling of PSMα2-3 derivatives.

Figure S4. Catalytic activity of PSMα2 and derivatives

Figure S5. Initial nitrocefin-degradation reaction rate in presence of PSMα3 and derivatives.

## AUTHOR INFORMATION

### Corresponding Author

Raz Jelinek

### Author Contributions

E.A, M.L and R.J designed the research; N.G contributed to the hypothesis and preliminary experiments; E.A. performed all the experiments involving PSMα peptides and derivatives (CD spectroscopy, FTIR spectroscopy, nitrocefin degradation kinetics, TEM sample preparation, fluorescent labelling and critical dissolution concentration). H.R, M.L and R.J supervised the research; E.A and R.J wrote the manuscript. All authors have given approval to the final version of the manuscript.

## ASSOCIATED CONTENT

**Figure S1.**
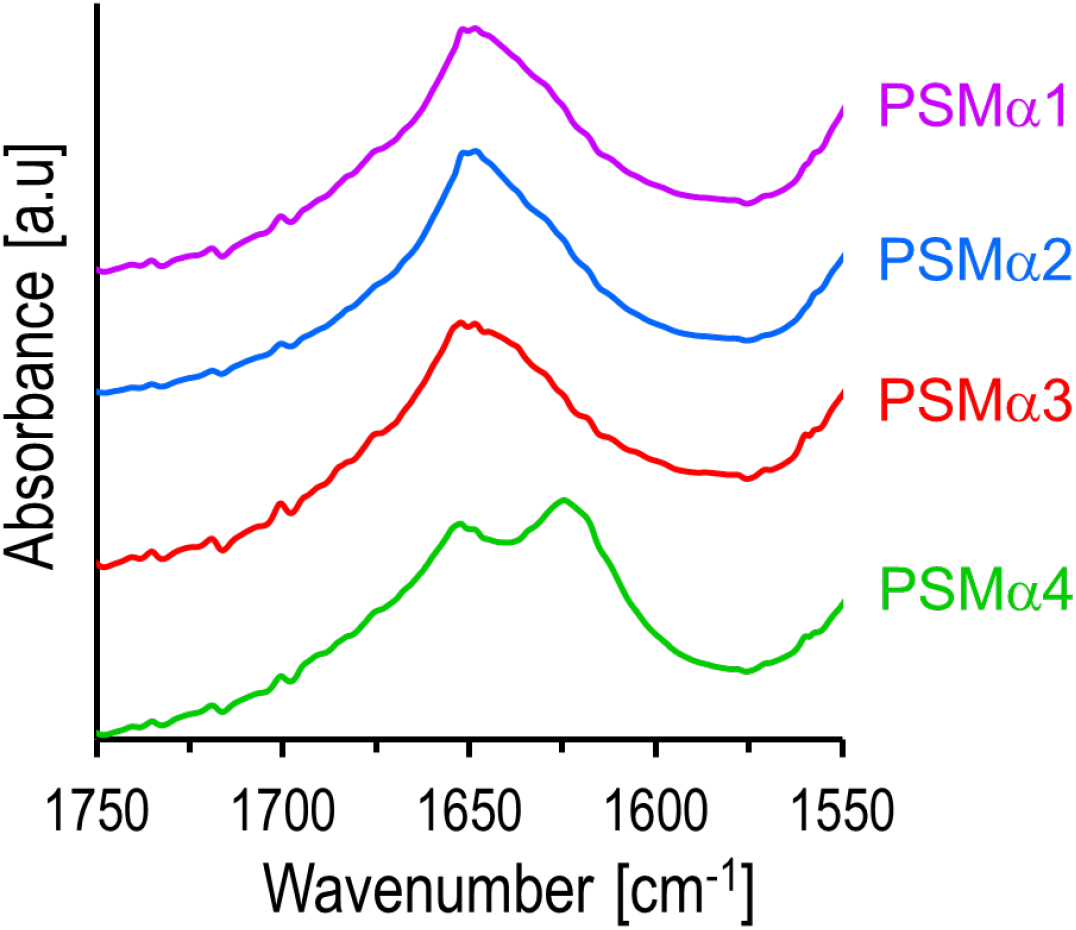
Fourier transform infrared spectroscopy of PSMα1-4.

**Figure S2.**
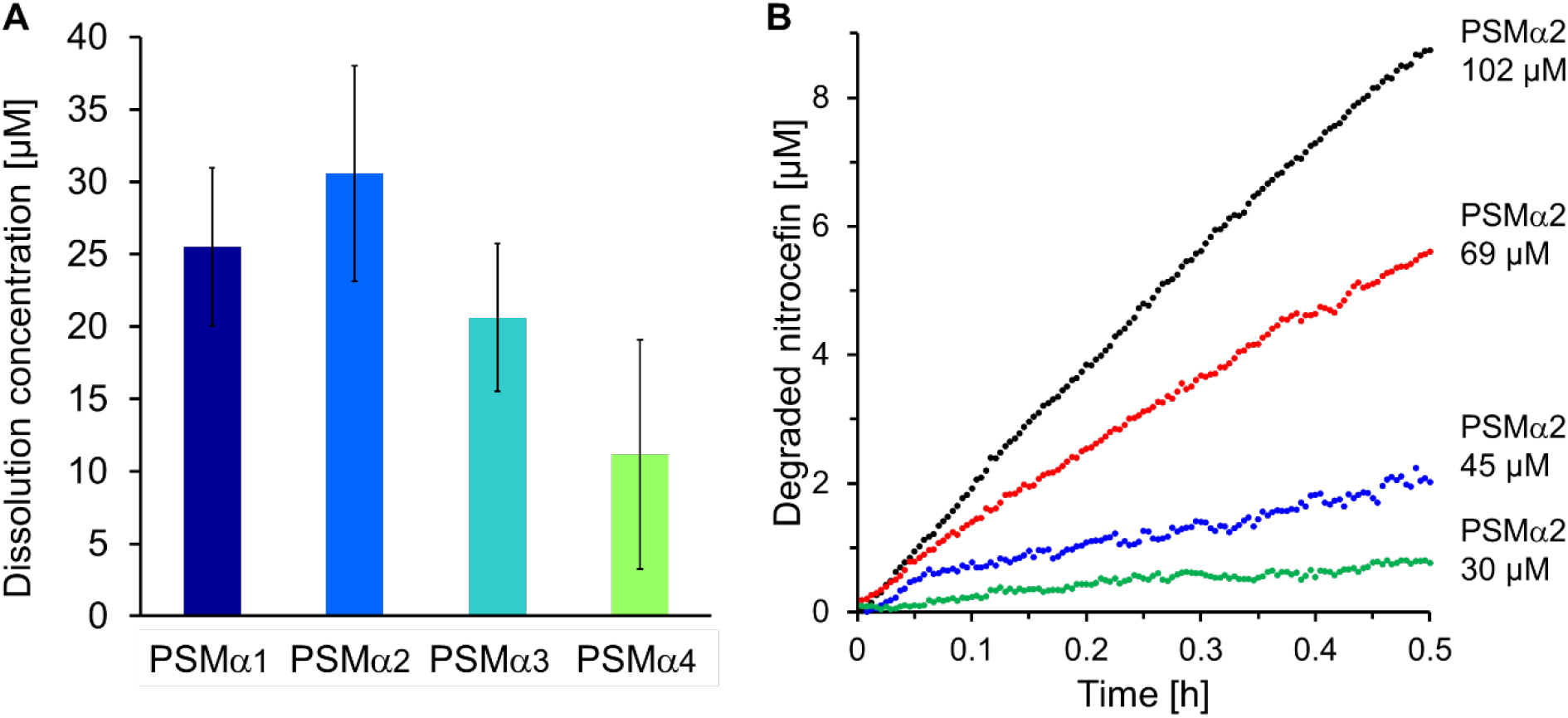
Critical dissolution concentration of PSMα1-4. **A**. The critical dissolution concentration was measured using ANS fluorescent labelling. The PSMα peptides were assembled as described above, incubated for two hours in DIW and buffered with HEPES at PSMα concentration of 440 μM. After the incubation the samples were diluted serially with the same buffer (HEPES 50 mM, pH 7.4) by 2/3 each time. PSMα solution (55 μL) was mixed with ANS aqueous solution (5 μL, 0.02 mg/mL) and placed in 384 well black plate. The samples were incubated for 30 minutes and then fluorescent measurement of ANS was applied (Excitation at 380 nm, emission 500 nm) at Biotek Synergy H1 plate reader (Biotek, Winooski, VT, USA). The fluorescence was analyzed as function of the peptide concentration (in logarithmic scale). The critical concentration was determined as the point in which the slope of the fluorescence increases dramatically (the intercept of two linear trends of fluorescence as function of the peptide concentration)^63^. **B**. Nitrocefin degradation in presence of PSMα2 (below and above the critical dissolution concentration, nitrocefin concentration 60 μM).

**Table S1.** Catalytic parameters of PSMα1-4 amyloid assemblies.

| | $R^2$ | $K_M$<br>[mM] | $V_{max}$<br>[μM/h] | $K_{cat}$<br>*$10^4$ [1/s] | $K_{cat}/K_M$<br>[$M^{-1} s^{-1}$] |
| --- | --- | --- | --- | --- | --- |
| PSMα1 | 0.95 | 0.2±0.2 | 60±30 | 0.9±0.5 | 0.4±0.2 |
| PSMα2 | 0.99 | 0.23±0.08 | 240±50 | 3.9±0.8 | 1.7±0.6 |
| PSMα3 | 0.96 | 0.08±0.03 | 100±20 | 1.6±0.3 | 2.1±0.7 |
| PSMα4 | 0.98 | 0.2±0.2 | 80±50 | 1.4±0.8 | 0.6±0.2 |

Values derive from the fitting of the initial rates to MM model and presented as *average ± confidence interval*. $K_M$ is the Michalis constant; $V_{max}$ is the maximal rate of reaction; $k_{cat}$ is the turnover number, and the catalytic efficiency is $\varepsilon=K_{cat}/K_M$ .

**Figure S3.**
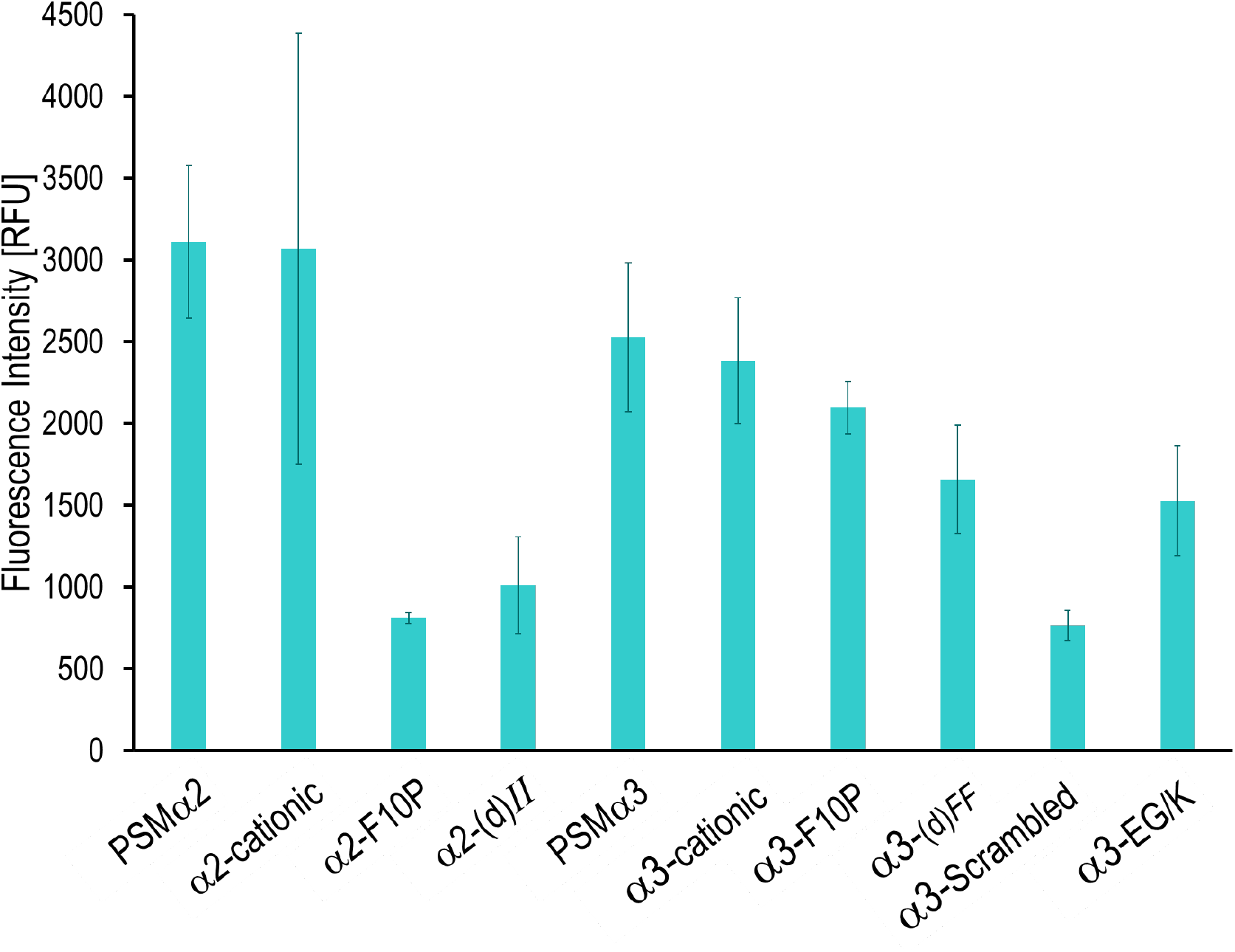
Amytracker-680 fluorescent labelling of PSMα2-3 derivatives. Amyloid staining with *Amytracker 680* (excitation 550 nm, emission 650 nm, PSMα concentrations were 400 μM).

**Figure S4.**
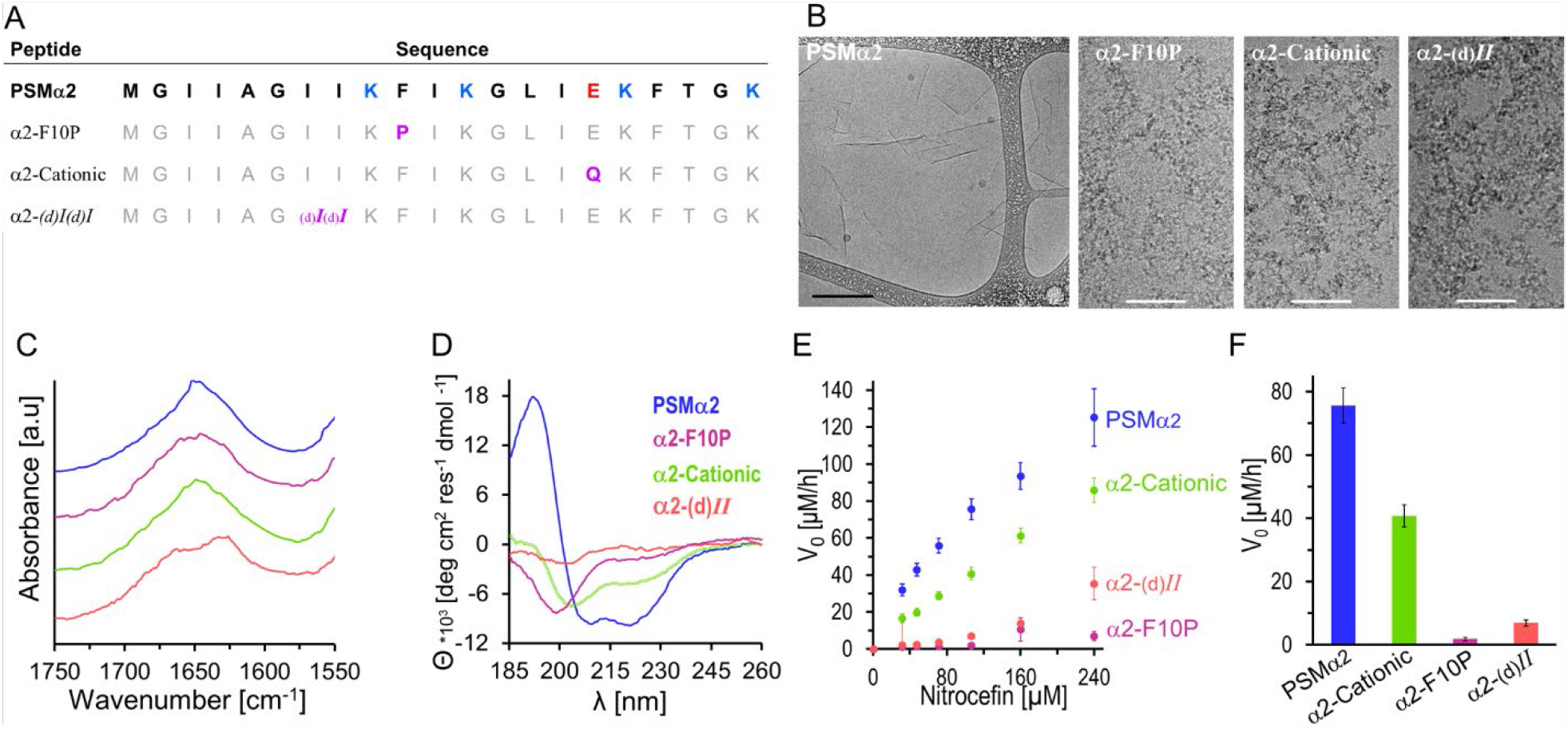
Catalytic activity of PSMα2 and derivatives. **A**. PSMα2 derivative sequences. The modified residues in the derivatives are in purple. At α2-cationic also the C-terminus is amidated. For α2-(d)*II* the two Ile residues are in (D) chirality rather than (L) like the other residues. **B**. Cryo-TEM images of PSMα2 derivatives (300μM). Only PSMα2 generated sheet morphology, while the others formed only amorphous structures. Black bar correspond to 500 nm and white bars correspond to 100 nm. **C**. Fourier transform infrared spectroscopy (FTIR) of amide vibration of PSMα2 derivatives. **D**. CD spectroscopy analysis of PSMα2 derivatives. **E**. Initial nitrocefin-degradation reaction rate, V_0_, as a function of initial substrate concentration in the presence of PSMα2 peptide assemblies (or derivatives, 170 μM). **F**. Initial degradation rate, V_0_, at initial nitrocefin concentration of 107 μM, in presence of PSMα2 and derivatives. The same color-coding was used for the PSMα2 derivatives.

**Figure S5.**
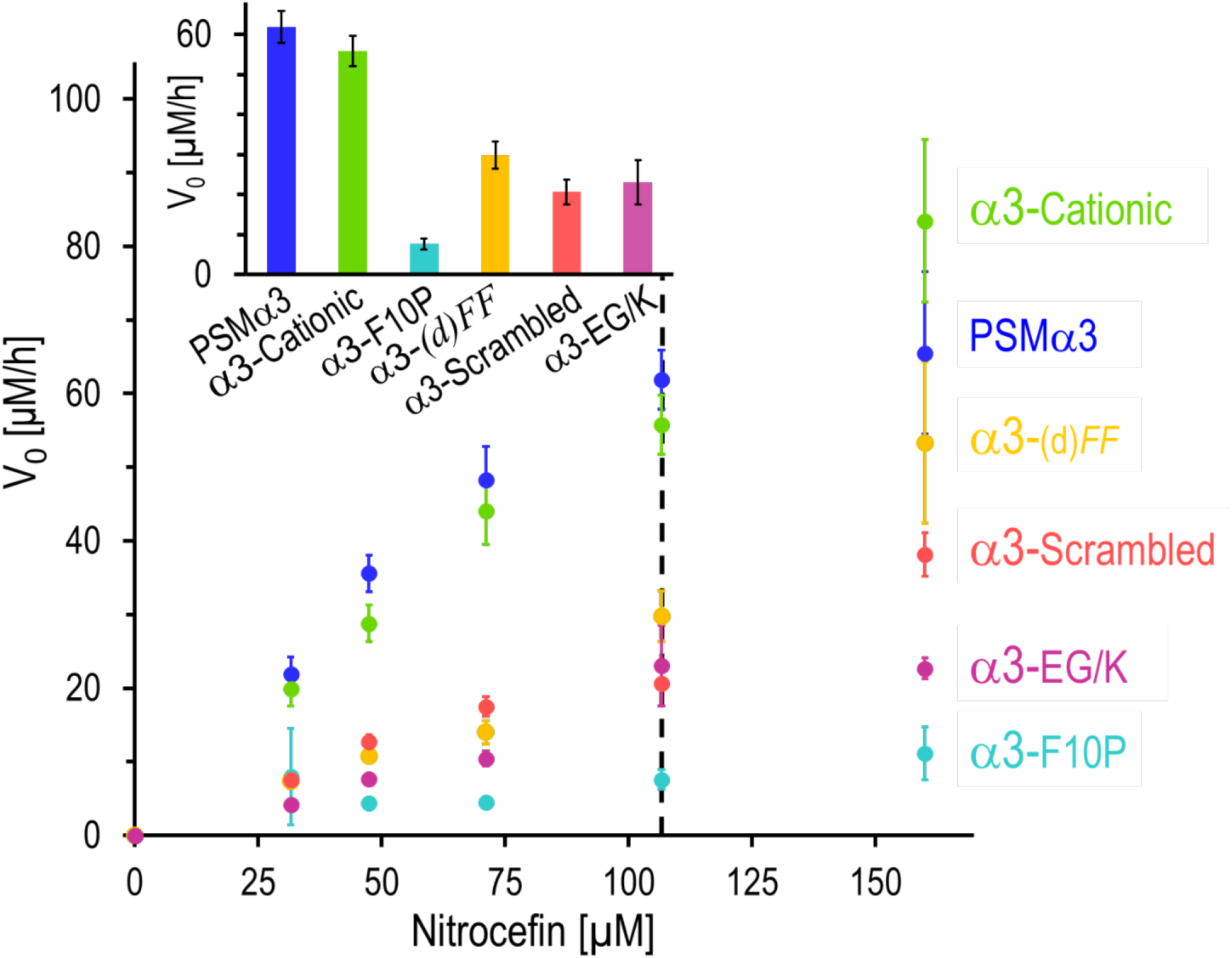
Initial nitrocefin-degradation reaction rate in presence of PSMα3 and derivatives. Initial nitrocefin-degradation reaction rate, V_0_, as a function of initial substrate concentration in the presence of PSMα3 peptide assemblies (or derivatives, 170 μM, broken line). At the inset: Initial degradation rate, V_0_, at initial nitrocefin concentration of 107 μM, in presence of PSMα3 and derivatives.

